# Mapping feather vane structure across the avian wing: spatial variation, asymmetry, and the effect of flight style

**DOI:** 10.64898/2026.06.08.730791

**Authors:** Gergely Osváth, Dragomir-Cosmin David, Dorottya Vargancsik, László-Jácint Nagy, Andrea Fehér, Zsolt Kovács, Ádám Zoltán Lendvai, Orsolya Vincze, Robert L. Nudds, Csongor I. Vágási, Péter L. Pap

**Affiliations:** Museum of Zoology, Babeș-Bolyai University, RO-400006 Cluj-Napoca, Clinicilor Street 5-7, Romania; Evolutionary Ecology Group, Hungarian Department of Biology and Ecology, Babeș-Bolyai University, RO-400006 Cluj-Napoca, Clinicilor Street 5-7, Romania; Department of Taxonomy and Ecology, Faculty of Biology and Geology, Babeș-Bolyai University, RO-400006 Cluj-Napoca, Clinicilor Street 5-7, Romania; Department of Evolutionary Zoology and Human Biology, University of Debrecen, H-4032 Debrecen, Egyetem sq. 1, Hungary; Department of Tisza Research, MTA Centre for Ecological Research-DRI, H-4026 Debrecen, Bem sq. 18/C, Hungary; School of Biological Sciences, Faculty of Biology, Medicine & Health, University of Manchester, Manchester, UK

**Author notes:** Correspondence: Gergely Osváth.

**Keywords:** barb density, barbule density, barb angle, feather vane asymmetry, flight style, vane macrostructure

## Abstract

Flight feather vanes are the primary aerodynamic surface of the avian wing. Because loading varies across the wing, vane macrostructure should co-vary with local mechanical demands, yet comparative data on how barb and barbule traits change among remiges and between vane surfaces remain scarce. We quantified barb density, barbule density, barb angle, barb length, and vane width on both vanes at three measurement positions along the rachis of all remiges in four species with contrasting flight modes (white stork, common buzzard, house sparrow, pygmy cormorant), generating over 40,000 measurements across 15 response variables from 992 feathers of 41 individuals. Two complementary generalised additive models characterised variation along the spanwise, inter-vane, and longitudinal axes, and compared outer primaries, inner primaries, and secondaries as functional wing regions. Feather macrostructure varied along all three axes and outer primaries represent the most distinctive region, with lower leading-vane barb density, reduced barb angles, and vane width asymmetry two to three times higher than in inner primaries or secondaries. House sparrow exhibited the densest vane architecture and the highest vane width asymmetry, whereas the low wing-beat frequency species showed complex nonlinear spanwise patterns undetectable by single-feather sampling. Pygmy cormorant barbule density was 39–53% lower than in all other species, matching its wettable plumage strategy. Longitudinal gradients in barb density and barb angle (22–31% decline) were conserved across species. The avian wing is thus functionally regionalised at the macrostructural level, with vane architecture reflecting both aerodynamic and ecological pressures.

**Summary statement:** Fine-scale vane measurements across all remiges in four species show macrostructural regionalisation of the avian wing, with outer primaries showing the most distinctive vane architecture.

## INTRODUCTION

Wing flight feathers (remiges) form the primary aerodynamic surfaces of the avian wing and play a fundamental role in flight ability and performance (Azuma, 2006; Norberg, 1990; Videler, 2005). A pennaceous flight feather consists of a central shaft (rachis) from which barbs extend laterally. Each barb bears distal and proximal barbules that interlock via a Velcro-like mechanism to form a continuous, planar vane surface (Bartels, 2003; Dyck, 2020; Stettenheim, 2000; Sullivan et al., 2017a; Tarsitano et al., 2000; Xu et al., 2001). The vane constitutes the main load-bearing and air-interacting component of the feather and is subject to complex aerodynamic forces during flight (Altshuler et al., 2015; Nafi et al., 2020; Tobalske, 2022; Usherwood, 2010). Therefore, the evolution of powered flight required morphological adaptations in the macrostructure of the remiges (Aparicio, 1998; Chang et al., 2019; Chuong et al., 2003; Lindhe Norberg, 2002; Norberg, 1985; Terrill and Shultz, 2023; Vágási et al., 2016).

During flight, the vane experiences two principal categories of mechanical loading. Out-of-plane forces are generated by the pressure differential between the dorsal and ventral wing surfaces, bend the vane perpendicular to its plane and are greatest during the downstroke when lift production peaks (Ennos et al., 1995; Usherwood, 2010). In-plane forces act within the vane surface, including shear along the barb axis and drag-induced tension at the leading edge where the airflow first contacts the feather (Ennos et al., 1995). The magnitude and temporal pattern of both force categories differ fundamentally among flight modes. Soaring and gliding species rely on external energy sources and experience relatively steady aerodynamic loads (Norberg, 1990; Pennycuick, 2008), whereas continuous flappers generate lift and thrust through sustained wing oscillations that impose cyclically varying forces on individual feathers throughout the wing-beat cycle. Passerine-type bounding flight introduces an additional dimension: brief, high-intensity flapping bursts alternate with ballistic pauses during which the wings are folded against the body, subjecting the remiges to abrupt transitions between loaded and unloaded states (Bruderer et al., 2010).

Aerodynamic loading is also non-uniform along three axes: (1) across the wing among different remiges; (2) across individual remiges between leading and trailing vanes; and (3) across individual remiges along the rachis from base to tip. Across the wing, distal feathers, which move at higher angular velocities during the wing stroke, experience greater aerodynamic forces per unit area than proximal feathers (Norberg, 1990; Usherwood, 2010). In many species, the outermost primaries separate during flight to form individual aerofoils at the slotted wing tip, each exposed to direct, unshielded airflow and concentrated loading that differs qualitatively from the shared load carried by overlapping inner primaries and secondaries (Tucker, 1993a; Videler, 2005). Within individual feathers, the narrow leading vane directly confronts the airflow and must resist bending and deformation, whereas the wider trailing vane, partially shielded by the overlap of adjacent feathers, primarily contributes to a continuous lifting surface and experiences different pressure distributions (Müller and Patone, 1998). Along the feather rachis, the base anchors the feather to the wing skeleton and must transmit the full aerodynamic load, while the tip is free to deflect under loading. These spatially varying force gradients predict that vane structure should differ systematically along all three axes (Bachmann et al., 2007; Crandell and Tobalske, 2011; Osváth et al., 2020; Pap et al., 2015a; Pap et al., 2015b; Pap et al., 2019).

The structural response of the vane to these forces is governed by multiple interacting traits. Barb density determines inter-barb spacing as a denser branching reduces gaps between adjacent barbs, lowering air transmissivity and enabling the vane to function as an aerodynamic surface more effective in force capture and transmission (Dial et al., 2012; Heers et al., 2011; Müller and Patone, 1998). Barbules maintain vane cohesion through a hook-and-groove interlocking mechanism between adjacent barbs; this adhesion increases structural robustness and delays yielding under repeated loading, and in case of eventual yielding vane integrity can be restored (Alibardi, 2007; Sullivan et al., 2016; Sullivan et al., 2017a). Barb branching angle influences vane rigidity by governing the alignment of barbs relative to the rachis with smaller angles increasing in-plane stiffness, characteristic of the narrow leading vane that directly confronts airflow (Feo et al., 2015; Norberg, 1985). Remiges’ vanes are asymmetric because the trailing vane is wider than the leading vane, and this asymmetry is determined by differences in barb length rather than in barb angle between the two vanes (Feo and Prum, 2014; Feo et al., 2015). Vane asymmetry contributes to feather rotation and torsional stability under aerodynamic load (Feduccia and Tordoff, 1979; Norberg, 1985; Prum and Williamson, 2001; Wang et al., 2019).

Despite the importance of vane structure for flight performance, integrative analyses remain limited. The clearest comparative signal of flight style comes from the most comprehensive phylogenetic analyses to date, those of Pap et al. (2015b, 2019), who examined up to 178 European bird species and found that soaring and gliding birds possess broader rachises but lower barb densities than active flappers, with barb angle on both inner and outer primaries differing among flight-type categories, although no consistent directional pattern emerged across groups (Pap et al., 2019). Feather structure also responded to aquatic habitat independently of flight type, body size, and wing morphology (Pap et al., 2015b), a result particularly relevant to interpreting the morphology of diving species such as cormorants.

How individual vane components contribute to mechanical behaviour has been addressed by a separate line of work. Ennos et al. (1995) showed experimentally in the pigeon (*Columba livia domestica*) that barb angle generates asymmetric resistance to dorsoventral loading, with vanes resisting forces from below more than from above, a bias that follows from the oblique attachment of barbs to the rachis; the same study found that outer primaries resist both out-of-plane and in-plane forces more than inner primaries and secondaries; although several feather types were examined, that study focused on mechanical behaviour and did not quantify systematic macrostructural variation across all flight feathers. Feo et al. (2015) subsequently established that barb angle covaries with vane function and that barb length, rather than barb angle, sets vane asymmetry across extant and fossil birds, although that work did not examine how these traits vary along the feather axis or among remiges across the wing, while Sullivan et al. (2016, 2017) characterised the barbule interlocking mechanism and found the branching pattern to be broadly conserved among flying birds. The functional weight of these traits is reinforced by Saitta et al. (2025), who found that barb angle asymmetry between vanes declines after independent evolutionary losses of flight, even as most other microscopic vane traits remain conserved, pointing to active flight selection on asymmetry against a developmentally constrained background.

Far less is known about how these traits are distributed within the wing and along individual feathers. Bachmann et al. (2007) provided the first systematic morphometric characterisation of wing feather macrostructure, measuring barb angle, barb density, and vane depth (equivalent to vane width in the present study) at 10% intervals along feathers of the barn owl (*Tyto alba*) and pigeon, and showed that the two species achieve a smooth vane surface through fundamentally different construction principles: the barn owl holds barb angle nearly constant along and across the wing, producing uniform in-plane stiffness, whereas the pigeon shows strong spanwise variation, so that vane rigidity is tuned differently under the contrasting loads of silent gliding and rapid flapping. The same spatial structuring is evident at finer scale within individual feathers and across the wing: barb density decreases along the rachis from base to tip, and outer primaries carry lower leading-vane barb density than inner primaries, while trailing-vane barb density shows the opposite pattern at the feather midpoint, indicating that the aerodynamic gradient along the wing shapes the two vane surfaces in opposite directions (Pap et al., 2019). These broad surveys, however, sampled only two feathers across the wing (one inner and one outer primary) and excluded secondary feathers, which form the main lifting surface of the inner wing and experience distinct aerodynamic loading; the recent finding that secondary vane dimensions scale with ulna length in an order-specific manner across 209 species (Deeming et al., 2025) further underscores the need to characterise secondary morphology in detail. To date, no study has integrated measurements of barb density, barbule density, barb angle, and vane width across all remiges while simultaneously resolving how these traits change along the base-to-tip axis of individual feathers and between leading and trailing vanes within the same individuals.

In this study, we investigate vane structural variation across four bird species representing distinct flight styles: the white stork (*Ciconia ciconia*; flapping-soaring), common buzzard (*Buteo buteo*; flapping-gliding), house sparrow (*Passer domesticus*; passerine-type flight), and pygmy cormorant (*Microcarbo pygmaeus*; continuous flapping). We quantify barb density, barbule density, barb angle, vane width, and their respective asymmetries to provide, to our knowledge, the first within-individual high-resolution mapping of vane macrostructure across both the longitudinal axis of individual feathers and the spanwise axis of the wing. Rather than maximising species number, we deliberately focus on a small set of species representing distinct flight styles, wing morphologies, and ecological strategies, enabling high-resolution within-wing and within-feather comparisons across species selected for contrasting flight styles, following an established approach for resolving fine-scale structural patterns before extending to broader phylogenetic sampling.

## MATERIALS AND METHODS

### Feather collection and measurements

Wing flight feathers were sampled from four bird species with contrasting flight styles: White stork, common buzzard, house sparrow, and pygmy cormorant. For each species, 10 individuals were examined, except for the white stork, where 11 individuals were sampled. Feather samples were collected from individuals which accidentally died due to various reasons. All remiges were fully grown and in good condition, with no visible signs of active moult or feather damage. White storks were either first-year birds, sub-adults, or adults; common buzzards and pygmy cormorants were first-year or adult birds; all house sparrows were in adult plumage. Regardless of age class, only individuals with fully developed flight feathers showing no signs of active growth were included. All primary and secondary feathers were collected from one wing of each individual. Detailed information on specimen origin and feather collection procedures is provided in Osváth et al. (2020).

The total number of sampled feathers differed among species due to interspecific differences in the number of remiges. In total, 32 feathers were collected from white storks (11 primaries and 21 secondaries), 24 from common buzzards (10 primaries and 14 secondaries), 22 from pygmy cormorants (10 primaries and 12 secondaries), and 18 from house sparrows (9 primaries and 9 secondaries). The innermost secondary (S22) in white storks was absent in all sampled individuals, while the outermost primary (P10) in house sparrows is vestigial; therefore, these feathers were excluded from the analysis.

Each feather was photographed on a metric grid background or next to a stage micrometer. Digital images were analysed using ImageJ (version 1.54f; National Institutes of Health, Bethesda, MD, USA; https://imagej.nih.gov/ij/). Spatial calibration was performed for each image by setting the known distance of the scale reference (metric grid or stage micrometer) in ImageJ, ensuring consistent µm/pixel resolution across all measurements. For each feather, measurements were taken separately on the leading and trailing vanes. Structural traits were quantified at three standardised measurement positions along the longitudinal axis of the vane: at 25% (base), 50% (middle), and 75% (tip) of the total vane length measured from the feather base (Fig. 1). In the house sparrows, measurements on the proximal secondaries S8 and S9 were restricted to two measurement positions (base and tip) due to the small size of these feathers.

**Figure 1.**
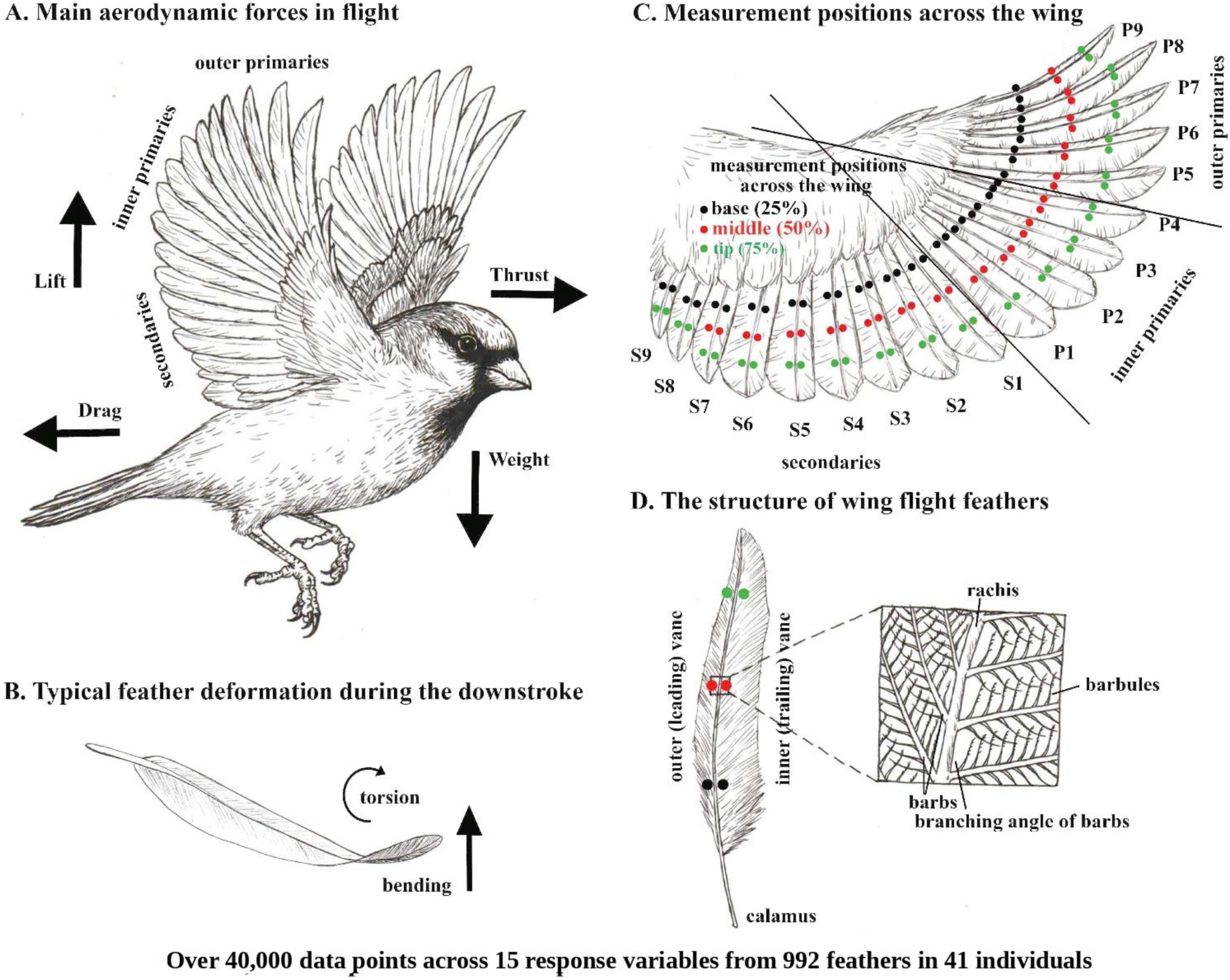
Schematic overview of aerodynamic forces, feather deformation, sampling design, and vane structure. (A) House sparrow in flight, illustrating the arrangement of outer primaries, inner primaries, and secondaries (remiges) and the four main aerodynamic forces acting during flight. (B) Typical feather deformation during the downstroke, showing bending perpendicular to the vane plane and torsional twisting along the rachis axis. (C) Dorsal view of the spread wing showing the sampling method, the three standardised measurement positions along the longitudinal axis of each feather vane: base (25%, black), middle (50%, red), and tip (75%, green). (D) Structure of a single wing feather showing the outer (leading) and inner (trailing) vanes, rachis, calamus, and the three measurement positions. An interactive version of the feather structure with all measured traits is available in Supplementary Material S1 (Feather Visualizer).

Barb density was quantified as the number of barbs per centimetre of rachis length. At each measurement position, barbs were counted within a 1-cm segment of rachis centred on the target measurement position, separately on the leading and trailing vanes, providing a single density value per vane at each measurement position.

Barbule density was quantified as the number of barbules per millimetre of barb length. Measurements were taken along a single, representative barb located closest to the target measurement position on the rachis. Barbules were counted on the distal side of the barb only, as distal barbules form the interlocking mechanism of the vane and are most directly involved in maintaining vane cohesion under aerodynamic loading (Sullivan et al., 2017a). This standardisation follows the functional role of distal barbules in vane integrity. Proximal barbules were not considered because they primarily function in overlap rather than mechanical coupling.

Barb branching angle was measured as the angle between the rachis and the barb, using the same representative barb selected for barbule density measurements. Barb length was measured as the total length of one barb per measurement position in millimetres.

Vane width was measured as the perpendicular distance between the rachis and the edge of the vane in a plane orthogonal to the rachis, separately for the leading and trailing vanes at each measurement position.

For each structural trait, vane asymmetry was calculated as the ratio of the trailing vane value to the corresponding leading-vane value at the same measurement position. To assess measurement reliability, a subset of feathers was re-measured, and repeatability was quantified using intraclass correlation coefficients (ICC). Repeatability estimates were high for all traits (barbule density: ICC = 0.875, 95% CI [0.77, 0.93]; barb density: ICC = 0.980, 95% CI [0.94, 0.99]; barb angle: ICC = 0.935, 95% CI [0.89, 0.96]; barb length: ICC = 0.994, 95% CI [0.98, 1.00]; vane width: ICC = 0.998, 95% CI [0.99, 1.00]; all p < 0.0001).

### Statistical analysis

All statistical analyses were performed using R version 4.4.3 (R Core Team, 2025). Feather vane morphology was analysed using generalised additive models (GAMs) implemented in the mgcv package (Wood, 2017). GAMs were chosen because they can flexibly model nonlinear variation in morphological traits across the wing (spanwise variation) while accounting for the factorial structure of the experimental design.

In what follows, two spatial axes are distinguished. The spanwise axis runs across the wing among feathers, from the outermost primary to the most proximal secondary, and is parameterised by the continuous variable span_pos and the categorical feather region factor. The longitudinal axis runs along the rachis of each individual feather, from base to tip, and is parameterised by the categorical *measurement_position* factor (three levels: base, middle, tip). Throughout, “position” refers exclusively to the longitudinal measurement position along the rachis.

We analysed 15 response variables organised into five trait categories: (1) barb density (n/cm) on leading and trailing vanes, plus asymmetry ratio; (2) barb angle (degrees) on leading and trailing vanes, plus asymmetry ratio; (3) barb length (mm) on leading and trailing vanes, plus asymmetry ratio; (4) barbule density (n/mm) on leading and trailing vanes, plus asymmetry ratio; and (5) vane width (mm) on leading and trailing vanes, plus asymmetry ratio. Asymmetry was calculated as trailing/leading ratios, where values >1 indicate larger values on the trailing vane. Ratios were used because they are scale-free and enable direct comparison across traits with different units and magnitudes.

To address two complementary questions about spatial variation in vane morphology, we employed two modelling approaches applied to the same dataset. Model 1 (spanwise smooth model) captures continuous nonlinear variation in each trait along the wing using species-specific smooth functions and quantifies how traits change along the base-to-tip axis of individual feathers. Model 2 (feather region model) formally tests differences among three functional wing regions (outer primaries, inner primaries, and secondaries) defined by species-specific breakpoints based on wing-tip slot morphology. Both models were fitted as generalised additive models (GAMs) using the mgcv package (Wood, 2017), with identical random-effect structures and the same 15 response variables.

Model 1 was specified as:

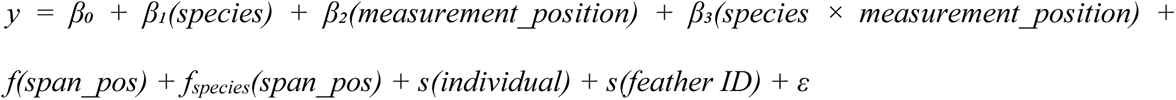

where β₀ is the intercept, β₁-β₃ are parametric coefficients for species (4 levels), measurement_position (3 levels: base, middle, tip), and their interaction, f() represents the overall smooth function for spanwise variation (span_pos), and f_species_() represents species-specific smooth deviations from the overall trend. The spanwise position variable (span_pos) was calculated within each species as a continuous variable ranging from 0 (most distal primary near the wingtip) to 1 (most proximal secondary near the body), standardising the spanwise axis across species with different numbers of flight feathers. Smooth terms were fitted using cubic regression splines (bs = “cr”) with basis dimension k = 10 for both the overall smooth and species-specific deviations (the latter with first-derivative penalty, m = 1). The factor-smooth interaction (by = species) allowed each species to have its own nonlinear pattern of spanwise variation while sharing the overall trend. Model 2 was specified as:

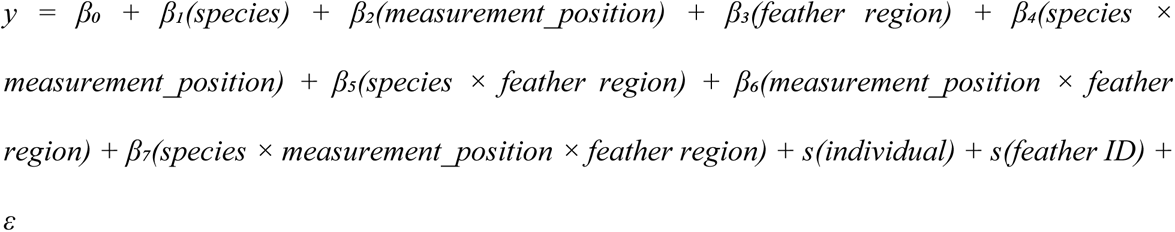

where feather region is a three-level factor (secondary, inner primary, outer primary) included with all two-way and three-way interactions with species and measurement_position (4 × 3 × 3 = 36 cells). The distinction between outer and inner primaries was based on species-specific breakpoints corresponding to the primaries that visibly separate at the slotted wing tip during flight: P1–P5 in common buzzard (10 primaries), P1–P6 in the white stork (11 primaries), P1–P4 in house sparrow (9 primaries), and P1–P4 in the pygmy cormorant (10 primaries), where P1 denotes the outermost primary. Model 2 did not include smooth terms for spanwise position, because the categorical feather region factor and the continuous span_pos variable are deterministically confounded (outer primaries always occupy the lowest span_pos values, secondaries the highest); including both would result in complete concurvity and unreliable parameter estimates. Instead, Model 2 treats the three wing regions as discrete functional units and tests for differences in trait means among them.

Both models included two crossed random intercepts to account for the hierarchical data structure: s(identity, bs = “re”) for individual-level variation and s(feather_id, bs = “re”) for feather-within-individual variation, where feather_id is a unique identifier for each individual × feather combination. These random effects partition variance attributable to among-individual and among-feather differences from the fixed effects of interest. Given the large sample sizes (n = 2,173–2,806 per response variable), models were fitted using bam() with fast restricted maximum likelihood (fREML) estimation and the discrete = TRUE option for computational efficiency (Wood, 2017), which provides equivalent results to standard REML for large datasets while substantially reducing computation time.

Estimated marginal means (EMMs) were calculated using the emmeans package (Lenth, 2025). For Model 1, EMMs were extracted for each species and measurement_position. For Model 2, EMMs were additionally extracted for each feather region and for the species × feather region interaction, enabling pairwise comparisons among all three wing regions both overall and within each species. Pairwise comparisons were performed with Tukey’s HSD adjustment for multiple comparisons. Effect sizes are reported as differences in EMMs with associated standard errors and 95% confidence intervals.

Model assumptions were verified by examining residual plots, Q-Q plots of deviance residuals, and the relationship between fitted values and residuals. For Model 1, basis dimension adequacy was confirmed using the k-index diagnostic (k.check in mgcv), which indicated that the chosen k = 10 provided sufficient flexibility for all smooth terms (all k-index values ≥ 1.0). The effective degrees of freedom (edf) of smooth terms were examined to assess the complexity of nonlinear relationships; edf ≈ 1 indicates a near-linear relationship, while higher values (edf > 3) indicate more complex, nonlinear curves. Model fit was assessed using adjusted R² and the percentage of deviance explained.

All analyses were conducted using the following R packages: mgcv v1.9-0 for GAM fitting via bam(), emmeans v1.10.7 for estimated marginal means and post-hoc comparisons, and multcomp for multiple comparison adjustments. Raw data and species-level patterns can be explored interactively in the Supplementary Material S1 (interactive data explorer). GAM smooth functions, spanwise variation plots, and estimated marginal means for all 15 response variables are provided in the Supplementary Material S2 (Figs S1–S5 and S16–S20). Full statistical tables, including model fit statistics, parametric coefficients, estimated marginal means, and pairwise comparisons, are provided in Supplementary Material S3 (Tables S1–S9).

### Ethical statement

All specimens used in this study were dead individuals obtained opportunistically (road kills, window strikes, or natural mortality). No live animals were handled or sacrificed for the purposes of this research; therefore, no ethics approval was required under current institutional and national regulations.

## RESULTS

### Model performance and variance partitioning

Two complementary GAMs were fitted to all 15 vane morphological traits. Model 1 (spanwise smooth) captured continuous nonlinear variation along the wing, explaining 62.1–97.2% of deviance for the primary structural traits (R²adj = 0.61–0.96; Table S1; smooth term details in Table S8). Model 2 (feather region) formally tested differences among three functional wing regions and showed comparable or slightly higher fit across all traits (72.6–97.8%; R²adj = 0.69–0.97), confirming that the three-level feather region factor captures a substantial portion of the spanwise variation. Asymmetry ratio models showed moderate fit in both models (51.2–72.7% in Model 1; 57.6–78.5% in Model 2). Both random intercepts contributed meaningfully to variance partitioning: the individual-level term was substantial in most models (edf = 17–35), while the feather-level term showed high effective degrees of freedom across most traits (edf = 202–728; the sole exception was vane width asymmetry, edf ≈ 0), indicating substantial among-feather variation within individuals.

Barb length and vane width showed strong allometric scaling with body size, with approximately 5- to 6-fold differences between the largest (white stork) and smallest (house sparrow) species (Table S9). Given this strong size-dependence, the following analysis focuses on the remaining 11 size-independent traits: barb density (leading, trailing, asymmetry), barbule density (leading, trailing, asymmetry), barb angle (leading, trailing, asymmetry), barb length asymmetry, and vane width asymmetry. Although barb length and vane width are size-dependent in their absolute values, their asymmetry ratios are size-independent; barb length asymmetry shows strong covariation with vane width asymmetry and is therefore not discussed in detail. Spanwise variation in barb length is illustrated in Fig. 5.

### Model 1: variation across the wing

Species-specific smooth terms were significant for the majority of traits, confirming that vane macrostructure varies nonlinearly along the wing in a species-dependent manner (Table 1; Figs 2–6). The overall smooth (s(span_pos)), representing the shared spanwise trend, was penalised to near zero (edf < 0.1) for approximately half of the size-independent traits (5 of 11), while the remaining traits showed low to moderate overall smooths (edf = 1.2–4.6), indicating that spanwise variation was predominantly species-specific but with a shared component for some traits.

**Figure 2.**
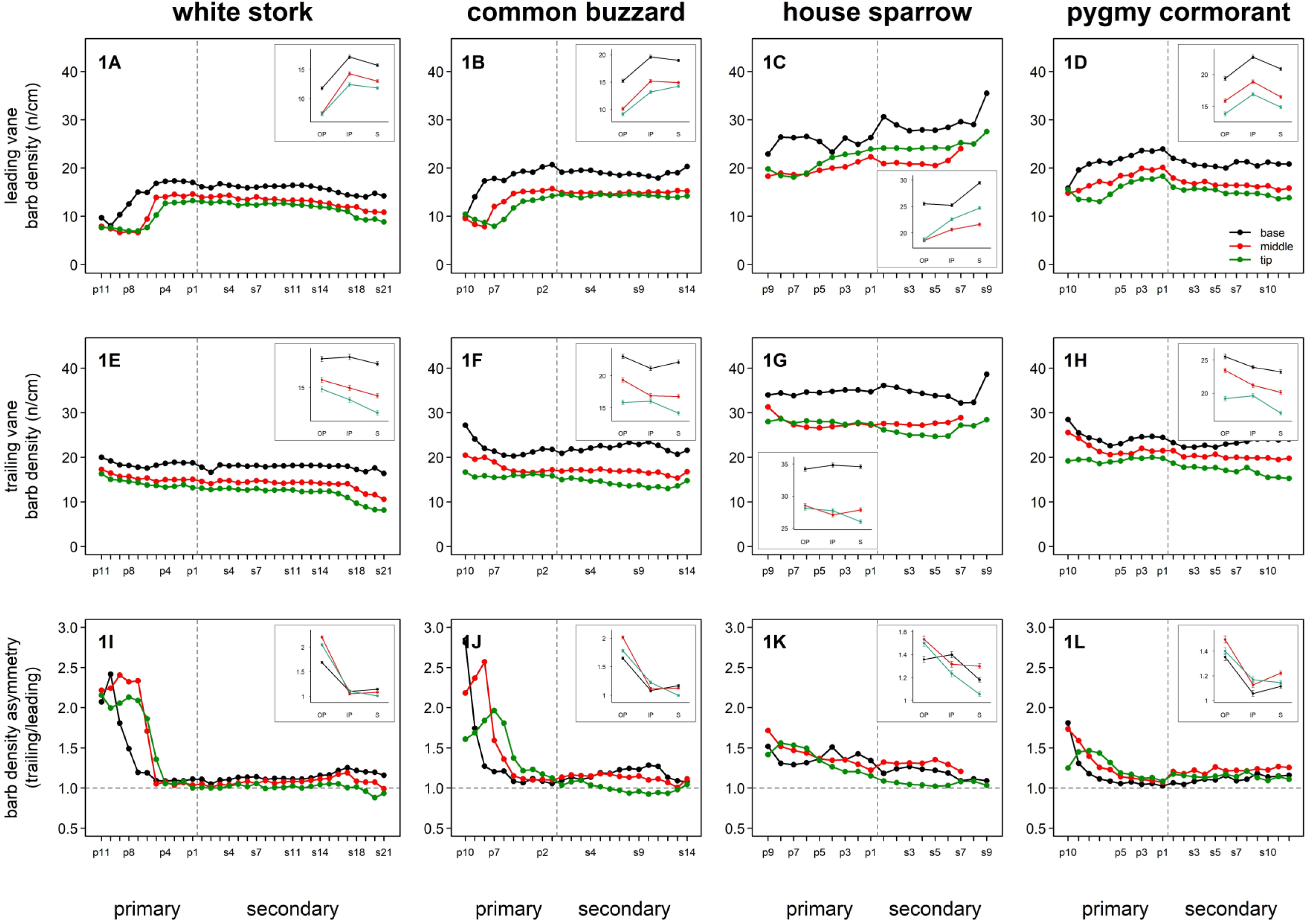
Barb density variation across the wing feather series in four avian species: White stork, common buzzard, house sparrow, and pygmy cormorant. Upper row (A–D): leading vane barb density (n/cm); middle row (E–H): trailing vane barb density (n/cm); lower row (I–L): barb density asymmetry (trailing/leading ratio). Within each panel, measurements are shown at three measurement positions along the vane: base (black), middle (red), and tip (green). Primary and secondary feathers are separated by a vertical dashed line. Error bars represent ± s.e.m. The horizontal dashed line in asymmetry panels indicates a ratio of 1.0 (symmetry). Each panel includes an inset showing estimated marginal means (± s.e.m.) from Model 2 for the three feather regions (OP = outer primary, IP = inner primary, S = secondary), with measurement positions colour-coded as in the main panels.

**Figure 3.**
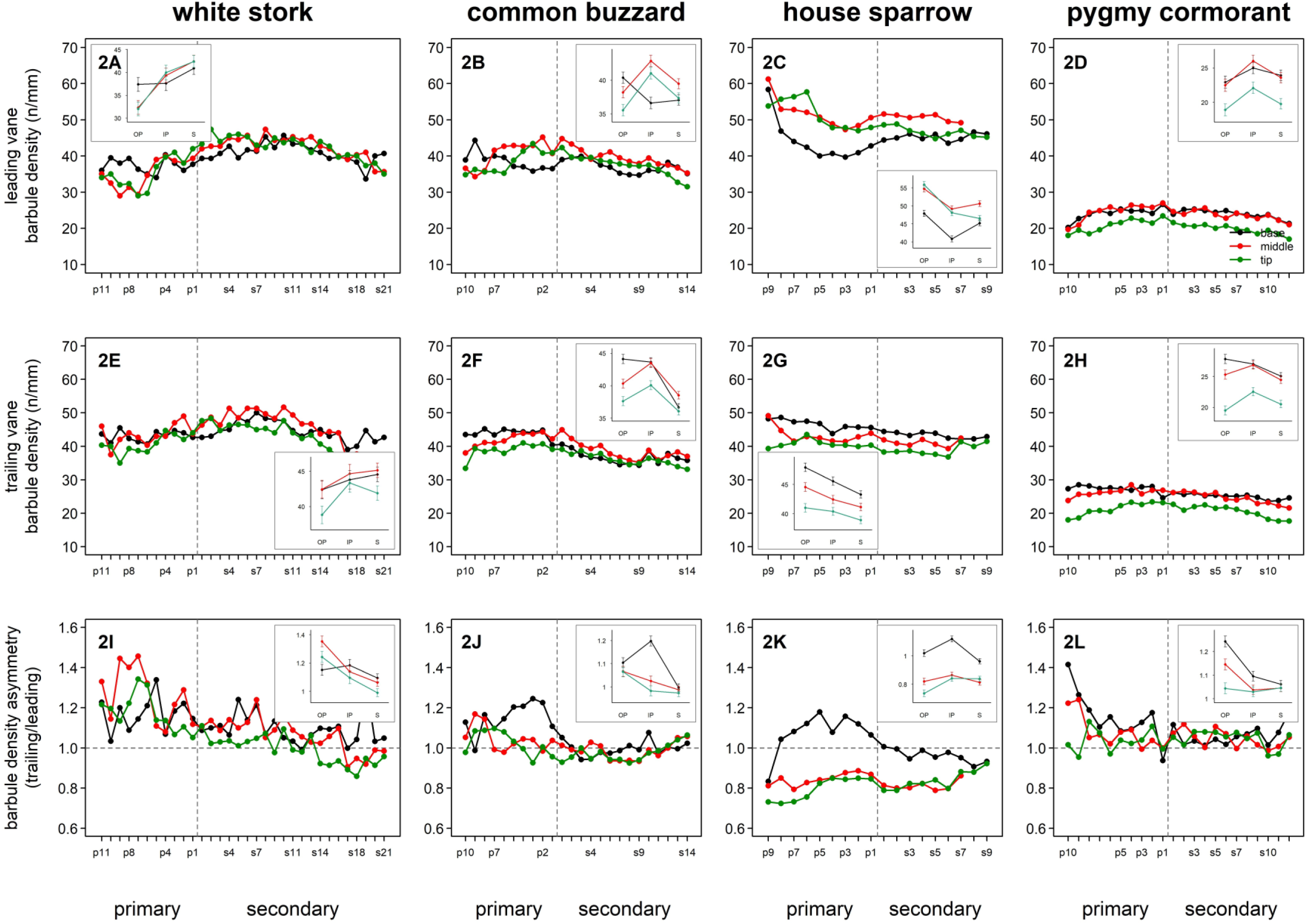
Barbule density variation across the wing feather series in four avian species. Upper row (3A-3D): leading vane barbule density (n/mm); middle row (3E-3H): trailing vane barbule density (n/mm); lower row (3I-3L): barbule density asymmetry (trailing/leading ratio). Within each panel, measurements are shown at three measurement positions: base (black), middle (red), and tip (green). Error bars represent ± s.e.m. The horizontal dashed line in asymmetry panels indicates a ratio of 1.0 (symmetry). Each panel includes an inset showing estimated marginal means (± s.e.m.) from Model 2 for the three feather regions (OP = outer primary, IP = inner primary, S = secondary), with measurement positions colour-coded as in the main panels.

**Figure 4.**
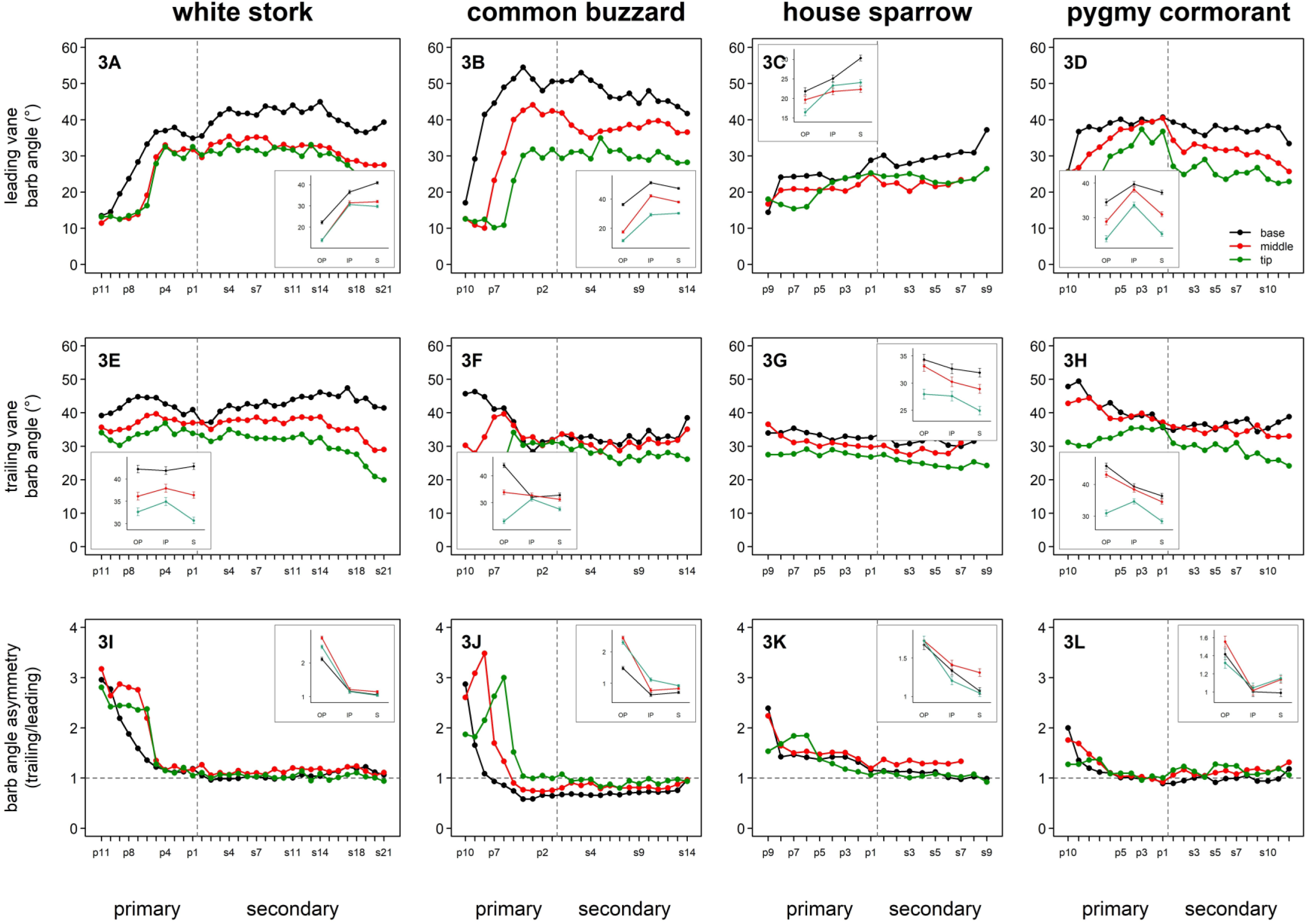
Barb angle variation across the wing feather series in four avian species. Upper row (4A-4D): leading vane barb angle (°); middle row (4E-4H): trailing vane barb angle (°); lower row (4I-4L): barb angle asymmetry (trailing/leading ratio). Within each panel, measurements are shown at three measurement positions: base (black), middle (red), and tip (green). Error bars represent ± s.e.m. The horizontal dashed line in asymmetry panels indicates a ratio of 1.0 (symmetry). Each panel includes an inset showing estimated marginal means (± s.e.m.) from Model 2 for the three feather regions (OP = outer primary, IP = inner primary, S = secondary), with measurement positions colour-coded as in the main panels.

**Table 1.** Effective degrees of freedom (edf) for smooth terms representing spanwise variation along the wing. Higher edf values indicate more nonlinear relationships. Significance: *** P < 0.001, ** P < 0.01, * P < 0.05, ns = not significant. ‘Overall’ = main spanwise trend across all species; species columns = species-specific deviation smooths (i.e. departures from the overall trend; the total spanwise curve for a given species is the sum of the overall and species-specific deviation terms). edf ≈ 1 indicates a near-linear but potentially strong effect; edf > 3 indicates complex nonlinear curves; edf ≈ 0 indicates no significant spanwise variation. Barb length and vane width smooth terms (leading and trailing) are omitted owing to strong allometric scaling (see Results); their asymmetry ratios are included.

| Trait | Overall | white stork | common buzzard | house sparrow | Pygmy Cormorant |
| --- | --- | --- | --- | --- | --- |
| Barb density (leading) | 0.00ns | 8.39*** | 8.58*** | 8.30*** | 7.28*** |
| Barb density (trailing) | 3.03ns | 7.84*** | 8.23*** | 5.69*** | 7.85*** |
| Barb density asymmetry | 2.63*** | 8.67*** | 8.79*** | 1.00*** | 5.21*** |
| Barbule density (leading) | 0.00ns | 6.71*** | 7.86*** | 5.89*** | 7.06*** |
| Barbule density (trailing) | 0.00ns | 7.96*** | 8.09*** | 2.63*** | 7.46*** |
| Barbule density asymmetry | 1.18ns | 3.01*** | 4.69*** | 3.74* | 5.08*** |
| Barb angle (leading) | 0.00ns | 8.59*** | 8.84*** | 4.18*** | 6.72*** |
| Barb angle (trailing) | 2.68* | 8.09*** | 7.77*** | 1.00*** | 6.91*** |
| Barb angle asymmetry | 3.37ns | 8.50*** | 8.78*** | 1.74*** | 2.32* |
| Barb length asymmetry | 0.00ns | 8.11*** | 7.70*** | 8.32*** | 6.82*** |
| Vane width asymmetry | 4.56*** | 6.78*** | 8.75*** | 7.89*** | 1.00*** |

The white stork and common buzzard exhibited the most complex spanwise patterns, with high effective degrees of freedom (edf = 3.0–8.7 and 4.7–8.8, respectively) for the species-specific smooths, indicating multiphasic nonlinear curves with local maxima and minima along the wing (Figs 2–6). The house sparrow showed a mixture of strongly nonlinear patterns (barb length asymmetry: edf = 8.3; barbule density: edf = 5.9; vane width asymmetry: edf = 7.9) and near-linear trends (barb density asymmetry, barb angle trailing: edf = 1.0). The pygmy cormorant showed significant species-specific smooths for all 11 size-independent traits, but with generally lower effective degrees of freedom (edf 1.0–7.9) and lower F-statistics than for the other species, pointing to more uniform macrostructural variation along its wing.

### Model 2: feather region differences

Outer primaries, inner primaries, and secondaries differed significantly in most macrostructural traits (Table 2; parametric coefficients in Table S2; pairwise comparisons in Table S4; visualised as estimated marginal means in the insets of Figs 2–6 and in detail in Figs S16–S20; interactive comparison in Supplementary Material S1, Region × Position). The largest differences were between outer primaries and the other two regions, while inner primaries and secondaries showed smaller or non-significant differences for several traits.

**Table 2.** Estimated marginal means (EMM ± s.e.m.) by feather region for size-independent vane traits (Model 2). Pairwise comparisons among feather regions (Tukey HSD). Secondary = reference level. Units: barb density (n/cm), barbule density (n/mm), barb angle (°). Asymmetry = trailing/leading ratio. Bold p-values indicate p < 0.001. S = secondary, IP = inner primary, OP = outer primary.

| Trait | Secondary | Inner primary | Outer primary | S vs IP | S vs OP | IP vs OP |
| --- | --- | --- | --- | --- | --- | --- |
| | EMM ± SE | EMM ± SE | EMM ± SE | p | p | p |
| Barb density (leading) | 18.07 ± 0.11 | 18.24 ± 0.12 | 14.43 ± 0.13 | 0.143 | <0.001 | <0.001 |
| Barb density (trailing) | 20.49 ± 0.13 | 21.31 ± 0.14 | 22.22 ± 0.15 | <0.001 | <0.001 | <0.001 |
| Barb density asymmetry | 1.13 ± 0.01 | 1.16 ± 0.01 | 1.67 ± 0.01 | <0.001 | <0.001 | <0.001 |
| Barbule density (leading) | 37.45 ± 0.45 | 37.39 ± 0.49 | 36.56 ± 0.49 | 0.972 | 0.002 | 0.024 |
| Barbule density (trailing) | 36.36 ± 0.35 | 38.66 ± 0.40 | 37.66 ± 0.40 | <0.001 | <0.001 | 0.006 |
| Barbule density asymmetry | 0.99 ± 0.01 | 1.05 ± 0.01 | 1.08 ± 0.01 | <0.001 | <0.001 | 0.006 |
| Barb angle (leading) | 32.36 ± 0.29 | 33.57 ± 0.34 | 21.69 ± 0.35 | <0.001 | <0.001 | <0.001 |
| Barb angle (trailing) | 32.25 ± 0.35 | 34.50 ± 0.38 | 35.61 ± 0.38 | <0.001 | <0.001 | <0.001 |
| Barb angle asymmetry | 1.04 ± 0.02 | 1.09 ± 0.02 | 1.91 ± 0.02 | 0.008 | <0.001 | <0.001 |
| Barb length asymmetry | 1.31 ± 0.03 | 1.88 ± 0.03 | 1.97 ± 0.04 | <0.001 | <0.001 | <0.001 |
| Vane width asymmetry | 1.78 ± 0.03 | 2.96 ± 0.04 | 5.16 ± 0.04 | <0.001 | <0.001 | <0.001 |

Barb density showed opposing vane-specific regional patterns. On the leading vane, outer primaries showed significantly lower barb density (EMM = 14.43 ± 0.13 n/cm) than both inner primaries (18.24 ± 0.12; p < 0.001) and secondaries (18.07 ± 0.11; p < 0.001), whereas inner primaries and secondaries did not differ (p = 0.143). On the trailing vane, the pattern was reversed: outer primaries showed the highest barb density (22.22 ± 0.15 n/cm), significantly exceeding both secondaries (20.49 ± 0.13; p < 0.001) and inner primaries (21.31 ± 0.14; p < 0.001). This opposing vane-specific pattern produced substantially higher barb density asymmetry in outer primaries (EMM = 1.67 ± 0.01), far exceeding inner primaries (1.16 ± 0.01) and secondaries (1.13 ± 0.01; both p < 0.001).

Barb angle showed a clear regional contrast on the leading vane. Outer primaries exhibited substantially lower leading-vane barb angles (EMM = 21.69 ± 0.35°) than inner primaries (33.57 ± 0.34°; p < 0.001) and secondaries (32.36 ± 0.29°; p < 0.001), reflecting the acute, cutting-edge morphology of wing-tip feathers. On the trailing vane, outer primaries showed the highest angles (35.61 ± 0.38°), slightly exceeding inner primaries (34.50 ± 0.38°; p < 0.001) and secondaries (32.25 ± 0.35°; p < 0.001). Barb angle asymmetry was, therefore, consequently highest in outer primaries (EMM = 1.91 ± 0.02), reflecting the large angular difference between leading and trailing vanes in the wing-tip region; inner primaries (1.09 ± 0.02) and secondaries (1.04 ± 0.02) showed near-unity ratios.

Barbule density showed a more moderate regional gradient. On the leading vane, the three regions did not differ substantially (secondaries: 37.45 ± 0.45; inner primaries: 37.39 ± 0.49; outer primaries: 36.56 ± 0.49 n/mm; outer vs. secondary p = 0.002; inner vs. secondary p = 0.972). On the trailing vane, inner primaries showed the highest barbule density (38.66 ± 0.40 n/mm), significantly exceeding secondaries (36.36 ± 0.35; p < 0.001) and outer primaries (37.66 ± 0.40; p = 0.006).

Vane width asymmetry showed the strongest regional differentiation among all traits examined. Outer primaries exhibited substantially elevated asymmetry (EMM = 5.16 ± 0.04)as compared with inner primaries (2.96 ± 0.04) and secondaries (1.78 ± 0.03; all p < 0.001). Inner primaries also showed significantly higher asymmetry than secondaries (p < 0.001), indicating a progressive increase in vane asymmetry from the wing root toward the wing tip.

Species × feather region interactions were significant for several traits (parametric coefficients in Tables S2, S4), indicating that the magnitude of differences among the three wing regions varied among species. These species-specific regional patterns are visible in the spanwise plots (Figs 2–6), where the transition between outer primaries, inner primaries, and secondaries corresponds to distinct inflection points in the curves.

### Interspecific variation in vane macrostructure

Species differed significantly in all macrostructural traits (Table S9; full pairwise comparisons in Table S6). House sparrow exhibited the highest barb density on both vanes, 1.3–1.9 times higher than that of the other species (Fig. 2). The pygmy cormorant showed distinctly reduced barbule density on both vanes, 39–53% lower than all other species (Fig. 3; all p < 0.001). The common buzzard exhibited the highest leading-vane barb angle (33.7°; Fig. 4). Pygmy cormorant showed the lowest (2.46), while House sparrow the highest vane width asymmetry (4.05; Fig. 6). Barb density asymmetry was above unity in all four species at the species level (range 1.23–1.38; Table S9), indicating consistently higher trailing-vane barb density. Among individual observation-level values, below-unity ratios occurred most frequently in common buzzard at the tip of the secondary feathers (Fig. 2).

### Longitudinal variation along the feather rachis

Macrostructural traits varied systematically along the feather rachis, with both consistent species-independent gradients and species × measurement_position interactions modulating the magnitude of these gradients (Figs 2–6; Table S3, Table S5, Table S7; interactive base/middle/tip profiles in Supplementary Material S1, Longitudinal profiles). Barb density decreased from base to tip on both vanes (leading: 20.15 → 14.99 n/cm, −26%; trailing: 24.82 → 18.68 n/cm, −25%; all contrasts p < 0.001). Barb angle showed a similar decrease (leading: 35.25° → 24.33°, −31%; trailing: 38.03° → 29.59°, −22%; all contrasts p < 0.001). These base-to-tip declines in both barb density and barb angle were consistent across all four species, indicating evolutionarily conserved flight-related structural gradients along the feather axis.

Barbule density showed contrasting longitudinal patterns between vanes. On the leading vane, barbule density peaked at the middle position (38.45 n/mm) relative to both base (36.31 n/mm) and tip (36.64 n/mm; base vs. middle and middle vs. tip p < 0.001; base vs. tip p = 0.204). On the trailing vane, barbule density decreased monotonically from base (39.33 n/mm) to tip (35.07 n/mm; all contrasts p < 0.001). Vane width asymmetry showed a significant longitudinal pattern, with higher values at the middle (3.39) and tip (3.38) positions than at the base (3.12; base vs. middle and base vs. tip p < 0.001; middle vs. tip p = 0.985).

Species × span_pos interactions were significant for several traits (Table S5), indicating that the magnitude of longitudinal change along the rachis varied among species in interaction with the spanwise location across the wing. The house sparrow showed particularly steep base-to-middle declines in barb density on both vanes relative to the other species (all p < 0.001), while species-specific longitudinal patterns in barbule density were more variable (Table S5). The magnitude of species differences also varied across measurement positions in a trait-specific manner. For barb angle on the leading vane, species differences were largest at the feather base (inter-species EMM range: 19.1°) and smallest at the tip (6.3°), indicating that the divergence in barb angle among species with distinct flight-styles is concentrated at the base of individual feathers. In contrast, species differences in barb density were similar in magnitude at the base and tip (leading vane: 11.9 vs. 11.5 n/cm), with species rankings preserved across measurement positions. For vane width asymmetry, the pattern was reversed: species differences were greater at the tip (range: 2.30) than at the base (0.91), driven primarily by elevated tip asymmetry in the house sparrow (Table S5).

## DISCUSSION

Our results show that feather vane macrostructure is organised along multiple spatial axes, with conserved longitudinal gradients along the rachis, marked inter-vane differences, and species-specific spanwise patterns that jointly define a functionally regionalised avian wing: outer primaries possess a distinctive vane architecture, characterised by reduced leading-vane barb density and barb angle, elevated trailing-vane barb density, and vane width asymmetry two to three times greater than in inner primaries or secondaries (Table 2). Among the traits examined, asymmetry ratios emerged as the most informative indicators of this regionalisation, with vane width asymmetry showing the largest and most consistent regional and interspecific differences. Although Bachmann et al. (2007) documented several vane traits at fine resolution in two species, to our knowledge no previous study has simultaneously characterised the variation of morphometric and geometric traits of the main structural components of the vane across all remiges across multiple species with contrasting flight styles at this resolution. This level of detail exceeds that of most previous comparative studies, which typically examined one or two feathers per individual (e.g., Pap et al. 2015b, 2019). The detailed within-wing and within-feather patterns documented here generate specific, testable hypotheses about functional morphology that complement existing works.

### Conserved within-feather gradients

Our three-point sampling along each feather’s rachis showed consistent longitudinal gradients that have not been systematically documented previously. The decrease in barb density (by 25–26%) and barb angle (by 22–31%) from base to tip observed across all four species likely reflects fundamental structural requirements for flight rather than flight-style-specific adaptations. The feather base must transmit aerodynamic forces to the wing skeleton and therefore requires maximum structural integrity, whereas the tip is free to deflect under load. Butler and Johnson (2004) found that barb breaking stress, strain, and toughness decrease distally along the rachis, demonstrating that longitudinal gradients in mechanical properties are a general feature of feather construction. That within-feather structural variation has functional consequences is further supported by de la Hera et al. (2020), who showed that European Robins (*Erithacus rubecula*) with longer migration distances develop primaries with shorter barbs and increased rachis thickness, resulting in stiffer feathers; this intraspecific evidence indicates that even modest differences in flight demands can shape the longitudinal distribution of structural elements within individual feathers. The evolutionarily conserved base-to-tip gradients across species with fundamentally different flight accords with the broader finding that most microscopic feather traits are developmentally constrained and evolve slowly even under relaxed selection following flight loss (Saitta et al., 2025), with the underlying developmental machinery sensitive to endocrine signals that modulate feather growth rate and final structural quality (Lendvai et al., 2021). These two patterns are reconcilable: a conserved structural foundation maintained by developmental constraints can still permit fine-scale adjustment under strong directional selection, as illustrated by the intraspecific differences documented in migratory passerines (de la Hera et al., 2020).

The non-monotonic pattern of barbule density on the leading vane, peaking at the middle position, was consistent across species and may represent a compensatory mechanism. The middle region of the feather may experience complex loading patterns during flight, including combined bending and torsional forces (Usherwood, 2010). Elevated barbule density in this region would enhance interlocking strength between barbs, thereby resisting separation forces under dynamic loading. This hypothesis could be tested through mechanical analysis of feather sections from different measurement positions along the rachis and possibly in multiple species with distinct flight-styles (Sullivan et al., 2016; Sullivan et al., 2017b).

### Spanwise variation and functional regionalisation

The fine-scale sampling across all flight feathers showed marked differences in spanwise variation among species (Table 1; Figs S1–S5 and S16–S20). The low wing-beat frequency species (white stork and common buzzard) exhibited the most conspicuous and complex multiphasic spanwise patterns. This indicates that the relatively steady but non-uniform aerodynamic loading during soaring and gliding drives regional specialisation of vane macrostructure along the wing (KleinHeerenbrink et al., 2017; Tucker, 1993b). Among the high wing-beat frequency species, house sparrow showed a mixture of strongly nonlinear and near-linear spanwise patterns across different traits, while the pygmy cormorant showed generally less complex almost linear pattern, as expected under more uniform aerodynamic loading during continuous flapping.

This dichotomy likely reflects different aerodynamic demands along the wing during different flight modes. In soaring and gliding flight, different wing sections experience distinct airflow regimes: the outer primaries function as individual aerofoils in slotted wing tips, whereas the inner wing provides the primary lifting surface (Videler, 2005). These spatially varying functional demands would favour regional specialisation of feather macrostructure. Pap et al. (2015b, 2019) demonstrated that feather structure differs between proximal and distal primaries in ways that correlate with flight mode, and Bachmann et al. (2007) documented variation among feathers across the wing in barb angle, barb density, and vane asymmetry, measured at 10% intervals along the rachis of barn owl and pigeon feathers, providing early evidence that spanwise gradients are a fundamental feature of flight feather functional morphology. Our high spatial resolution mapping extends these findings by showing that the pattern of variation along the wing is itself species-specific and more complex than a simple outer-to-inner gradient (Supplementary Material S1, Species × Region); the high edf values indicate multiphasic curves with local maxima and minima that may correspond to distinct functional regions.

The uniformity of the pygmy cormorant feather macrostructure across the wingspan is intriguing, and likely reflects the combined influence of two distinct selection pressures acting on the plumage of this foot-propelled diver: the aerodynamic demands of continuous flapping flight and the structural requirements of an aquatic lifestyle. During continuous flapping flight, the entire wing moves through similar aerodynamic regimes during each wing-beat cycle, in contrast to soaring flight where the outer primaries function as separated aerofoils in slotted wing tips while the inner wing provides the primary lifting surface (Videler, 2005). This interpretation is supported by Pap et al. (2015b), who reported that barbule density was highest in aquatic species with waterproof plumage; however, the apparent discrepancy with our results may reflect fundamental differences between wetting-resistant and wetting-tolerant aquatic birds.

Cormorants, in contrast to most aquatic birds, have a wettable plumage, an adaptation that facilitates diving by reducing buoyancy and eliminating air trapped within feathers (Grémillet et al., 2005; Rijke, 1968). Srinivasan et al. (2014) quantified feather wettability in six aquatic bird species, including three cormorant-family species, and showed that a dimensionless spacing ratio (D*) derived from contact angle measurements predicts the resistance of feathers to liquid penetration, with breakthrough pressures corresponding to depths of only 1–4 m. The reduced barbule density we observed in the pygmy cormorant matches this wettable plumage strategy: fewer barbules result in wider inter-barbule spacing, which increases D* and lowers breakthrough pressure, yielding a more porous and permeable vane structure that may facilitate water penetration at shallow dive depths. Nevertheless, direct measurements of feather permeability would be required to test this interpretation. Comparative works across diverse waterbird taxa generally identify aquatic ecology as a significant driver of feather macrostructure (Osváth et al., 2018; Pap et al., 2017; Pap et al., 2020).

Barbule density asymmetry showed a distinctive pattern in house sparrow: it was the only species with a ratio below 1.0 (0.89 ± 0.01), indicating higher barbule density on the leading than on the trailing vane. This asymmetry in barbule density possibly reflects the demand for robust leading-edge cohesion during rapid wing oscillations characteristic of passerine-type flight. By contrast, in the white stork and common buzzard, the trailing vane showed higher barbule density than the leading vane, matching the greater overlap and interlocking requirements of trailing vanes that form the main lifting surface during flight (Feo et al., 2015). The pygmy cormorant showed a nearly symmetric barbule density distribution across vanes (ratio = 1.08 ± 0.01), reflecting its uniformly reduced barbule density overriding any aerodynamic asymmetry signal.

The regional pattern of trailing-vane barbule density, where inner primaries showed the highest values (38.66 n/mm), significantly exceeding both secondaries (36.36) and outer primaries (37.66), was the only trait for which inner primaries surpassed outer primaries. This pattern may reflect the particular mechanical demands on inner primary trailing vanes, which form the continuous lifting surface of the mid-wing region and overlap extensively with adjacent feathers during flight (Matloff et al., 2020). Elevated barbule density in this region would strengthen inter-barb interlocking, thereby maintaining vane cohesion under the combined bending and shear forces that act on the overlapping inner wing surface during both steady flight and transitional manoeuvres (Sullivan et al., 2016; Sullivan et al., 2017a).

### Barb angle and vane rigidity

On the leading vane, both low wing-beat frequency species showed higher barb angles (species-level EMMs: 27.9–33.7°) than the house sparrow (22.8°; Fig. 4). This pattern aligns with the findings of Feo et al. (2015), who demonstrated that barb angle varies with vane function, and with broader phylogenetic findings on flight type and barb angle (Pap et al., 2019). Out-of-plane resistance and its directional asymmetry, however, are primarily governed by the three-dimensional inclination angle of barbs at their attachment to the rachis rather than by the in-plane branching angle measured here (Ennos et al., 1995). Bachmann et al. (2007) found that the barn owl maintains nearly constant barb angles along its feathers, in contrast to the pigeon, which shows substantial spanwise variation. The common buzzard exhibited the highest leading-vane barb angles (33.7°) among our species and showed the steepest base-to-tip decline in leading-vane barb angle (−47%), indicating that high absolute barb angle does not necessarily entail longitudinal uniformity across species with gliding-based flight styles.

The lower barb angles observed in the house sparrow may be associated with the dynamic wing shape changes required during flapping flight (Feo et al., 2015). During the upstroke, flight feathers rotate and separate to reduce air resistance, whereas during the downstroke they form a unified surface for thrust generation (Dvořák, 2016).

The white stork showed the largest barb angle asymmetry among the four species (EMM = 1.57), followed by the house sparrow (1.39), common buzzard (1.25), and pygmy cormorant (1.18). In all species, the trailing-vane barb angle exceeded the leading-vane angle, consistent with the more acute cutting-edge morphology of the leading vane. This pattern may relate to the slotted wing tip morphology characteristic of many soaring raptors, where individual primaries function as separated aerofoils (Tucker, 1993a). The functional relevance of barb angle asymmetry to flight is further supported by the finding that this trait shows the strongest and most consistent decrease following independent losses of flight across avian lineages (Saitta et al., 2025), implying that the interspecific variation we observe among our four volant species is maintained by active selection linked to aerodynamic function.

### Vane width asymmetry and species-specific profiles

Kiat and O’Connor (2024) demonstrated that vane asymmetry is functionally constrained in flying taxa: vane asymmetry shows distinct rates of change following independent losses of flight, underscoring that the degree of asymmetry is actively maintained by aerodynamic selection. Hall (2014) proposed that vane width ratios (termed “vane depth ratios” therein) exceeding 4:1 are required for feathers to exhibit stabilising torsional behaviour under aerodynamic load. The elevated asymmetry in the house sparrow compared to the low wing-beat frequency species in this study is noteworthy; however, whether this pattern reflects a general adaptation to fast flapping flight or is specific to house sparrow (or passerines) should be further explored. The importance of vane asymmetry as a flight adaptation is illustrated by comparative evidence from flightless birds: Saitta et al. (2025) showed that asymmetry-related traits exhibit the most prominent shifts following flight loss, whereas other vane properties remain relatively conserved over macroevolutionary timescales.

Beyond these species-level differences, the spanwise asymmetry profiles vary among feathers across the wing (Figs 5, 6; Figs S1–S5 and S16–S20). In the white stork and common buzzard, vane width asymmetry shows clear peaks at distinct locations along the wing, with the location of peak asymmetry varying among base, middle, and tip measurement points along the rachis. This peak structure indicates that the interaction between spanwise aerodynamic loading and longitudinal force gradients produces regionally specific vane proportions in these broad-winged species. In house sparrow, asymmetry peaks appear primarily at feather tips, reflecting the concentration of aerodynamic forces on the distal vane portion during high-frequency flapping. The pygmy cormorant, by contrast, exhibits comparatively flat asymmetry profiles across the wing, with minimal peak differentiation, paralleling its generally uniform macrostructural variation along the wing.

**Figure 5.**
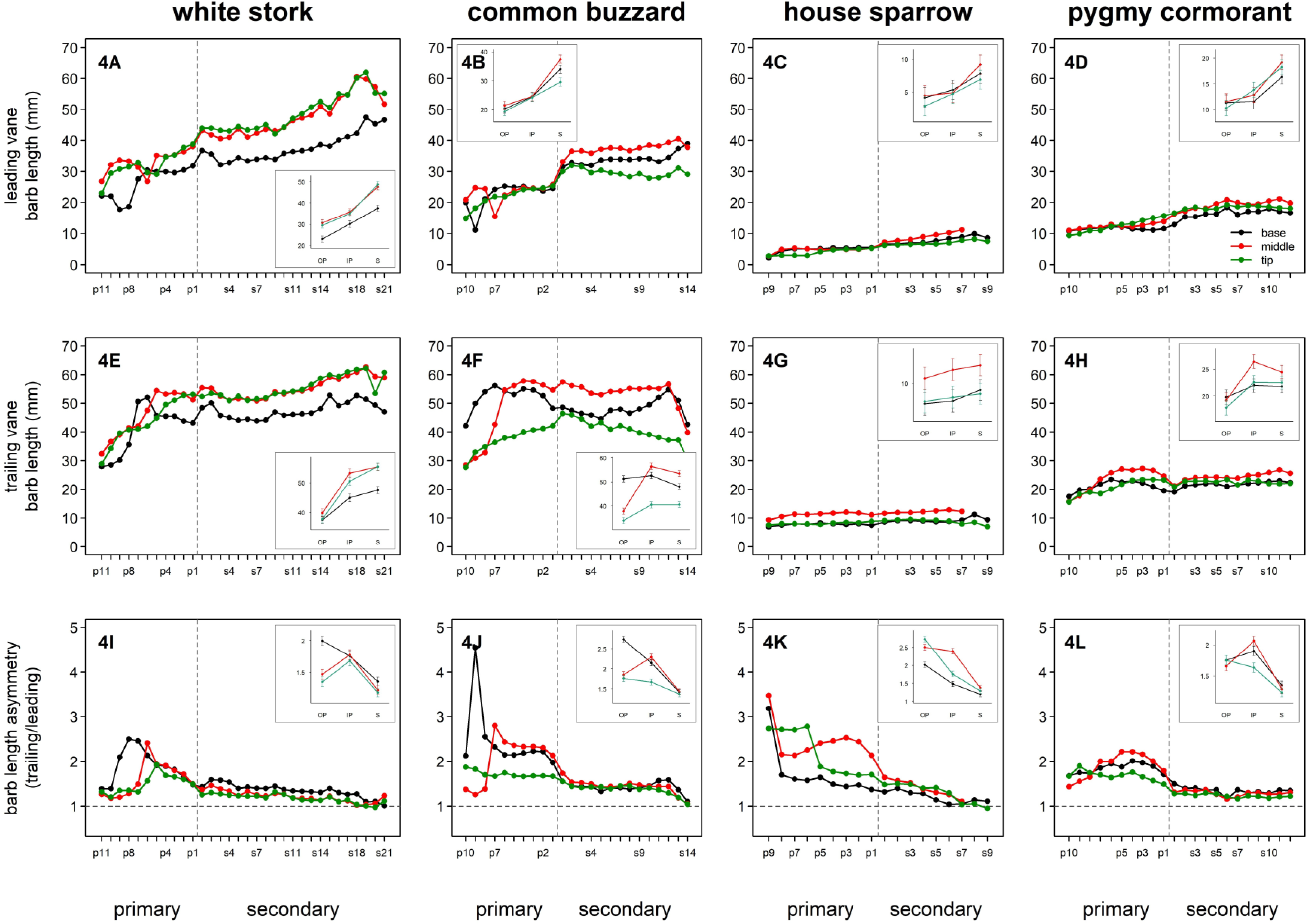
Barb length variation across the wing feather series in four avian species: White stork, common buzzard, house sparrow, and pygmy cormorant. Upper row (5A-5D): leading vane barb length (mm); middle row (5E-5H): trailing vane barb length (mm); lower row (5I-5L): barb length asymmetry (trailing/leading ratio). Within each panel, measurements are shown at three measurement positions along the vane: base (black), middle (red), and tip (green). Primary and secondary feathers are separated by a vertical dashed line. Error bars represent ± s.e.m. The horizontal dashed line in asymmetry panels indicates a ratio of 1.0 (symmetry). Each panel includes an inset showing estimated marginal means (± s.e.m.) from Model 2 for the three feather regions (OP = outer primary, IP = inner primary, S = secondary), with measurement positions colour-coded as in the main panels.

**Figure 6.**
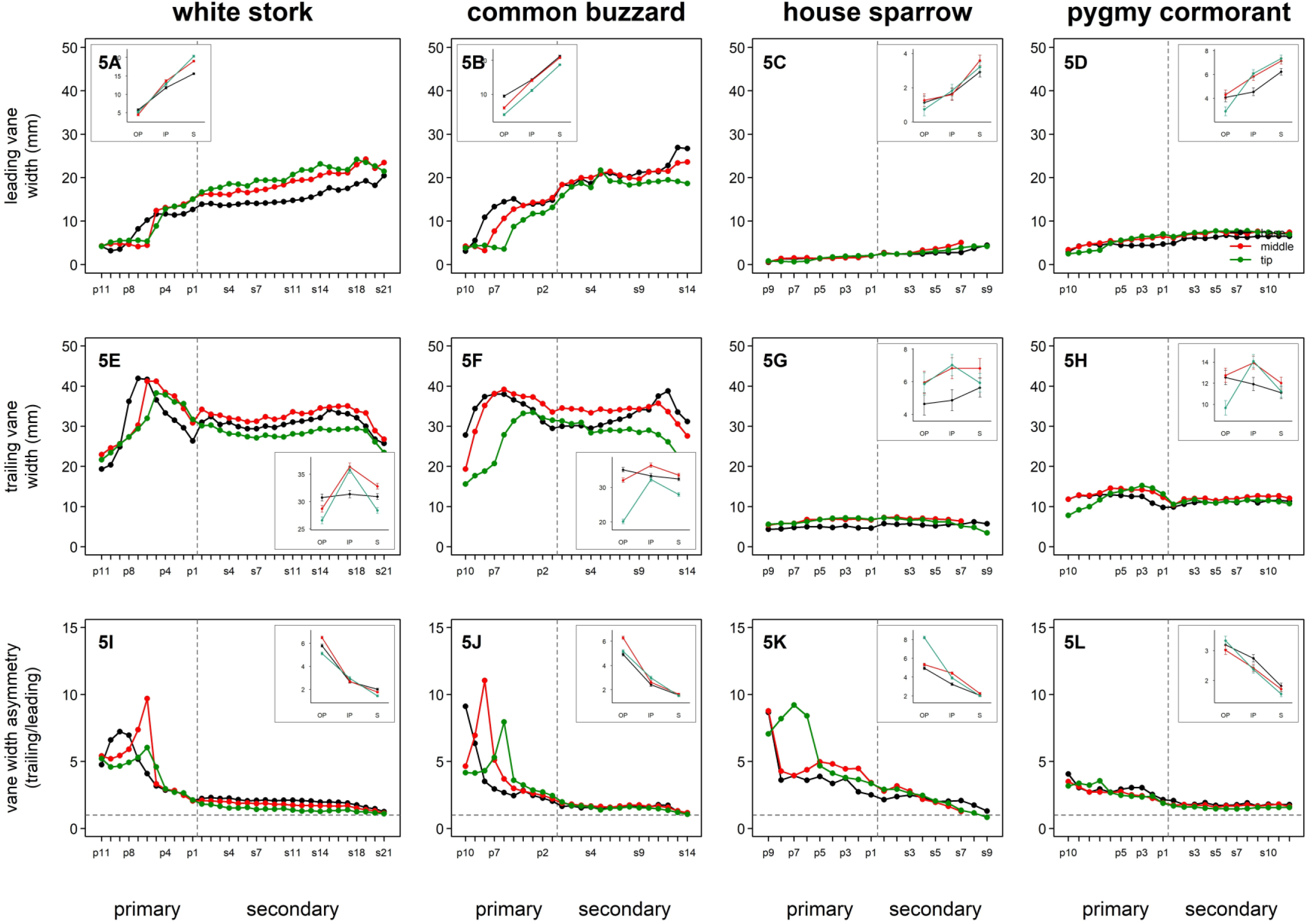
Vane width variation across the wing feather series in four avian species. Upper row (6A-6D): leading vane width (mm); middle row (6E-6H): trailing vane width (mm); lower row (6I-6L): vane width asymmetry (trailing/leading ratio). Within each panel, measurements are shown at three measurement positions: base (black), middle (red), and tip (green). Error bars represent ± s.e.m. The horizontal dashed line in asymmetry panels indicates a ratio of 1.0 (symmetry). Each panel includes an inset showing estimated marginal means (± s.e.m.) from Model 2 for the three feather regions (OP = outer primary, IP = inner primary, S = secondary), with measurement positions colour-coded as in the main panels.

The species-specific diagnostic profiles summarised in Table 3 (and visualised as multi-trait fingerprints in Supplementary Material S1, Species radar) integrate these trait-level patterns into coherent vane architectures for each species. The species ranking by vane width asymmetry (house sparrow 4.05× > white stork 3.45× > common buzzard 3.23× > pygmy cormorant 2.46×) indicates that this trait responds to multiple selection pressures. The elevated asymmetry in house sparrow likely reflects the demands of rapid, high-amplitude wing-beat kinematics, whereas pygmy cormorant’s low asymmetry parallels its reduced barbule density and may reflect the combined influence of aquatic ecology and wettable plumage strategy on vane proportions. Common buzzard’s lower asymmetry compared to white stork suggests that flapping-gliding and flapping-soaring flight impose different structural demands, though these species-level patterns cannot be disentangled from phylogenetic and allometric effects without broader taxonomic sampling.

**Table 3.** Species-specific diagnostic profiles of feather vane macrostructure. Summary of qualitative patterns across major structural traits measured in this study, including barb density, barbule density, barb angle, barb angle asymmetry, vane width asymmetry, and spatial variation along three measurement axes: the longitudinal axis (along individual feathers from base to tip), the spanwise axis (across the wing from distal primaries to proximal secondaries (spanwise variation among feathers)), and the inter-vane axis (between leading and trailing vanes). Descriptors (low, moderate, high) are relative to the four-species dataset and summarise dominant trends rather than absolute values. Functional interpretations synthesise structure-function relationships discussed in the main text and are based on established biomechanical and aerodynamic principles documented in previous studies (e.g. Ennos, 1995; Feo et al., 2015; Hall, 2014; Müller and Patone, 1998; Pap et al., 2015a; Pap et al., 2015b; Saitta et al., 2025; Srinivasan et al., 2014; Sullivan et al., 2016; Sullivan et al., 2017; Videler, 2005). Interpretations are inference-based and represent mechanistic associations between observed structural patterns and previously established biomechanical relationships rather than direct performance measurements. Trait values are estimated marginal means from Model 2 (Table S9), averaged across feather regions and measurement positions.

| Trait values |  |  |  |  |  | Spatial patterns |  |  |  |
| --- | --- | --- | --- | --- | --- | --- | --- | --- | --- |
| Species | Barb density (BD) | Barbule density (BbD) | Barb angle (leading vane) (BA) | Barb angle asymmetry (BAA) | Vane width asymmetry (VWA) | Longitudinal (base → tip) | Spanwise (across wing) | Inter-vane | Functional interpretation |
| **white stork** | **Low** (12.3 / 15.6) | **Moderate-high** (38.3 / 43.0) | **Intermediate** (27.9°) | **High** (1.57; T > L) | **High** (3.45×) | Clear base-to-tip decrease in BD (−29% / −27%) and BA (−25% / −23%); barbule density stable on leading vane, moderate trailing decline (−5%) | Strongly nonlinear for all 11 traits (edf 3.0-8.7); regionally structured with local maxima/minima | Moderate barbule asymmetry (1.15; T > L); trailing vane wider (3.45×); BD higher on trailing vane (1.38×) | *Reflects* stable vane geometry under steady soaring loads; complex spanwise variation indicates regional specialization matching heterogeneous airflow along the wing |
| **common buzzard** | **Low-intermediate** (14.5 / 18.4) | **Moderate** (38.7 / 40.1) | **Highest** (33.7°) | **Very high** (1.25; T > L) | **Very low** (3.23×) | Steepest leading BA decline (−47%) among species; moderate BD decline (−32% / −31%); barbule density peaks at middle on leading vane, moderate trailing decline (−9%) | Strongly nonlinear for all 11 traits (edf 4.7-8.8); highest minimum edf among species (4.7) | Weak barbule asymmetry (1.04; T ≈ L); BD higher on trailing vane (1.35×); second lowest VW asymmetry (3.23×) | *Reflects* stable, moderately asymmetric vane suited to gliding and slotted-tip aerodynamics; steep base-to-tip BA gradient on leading vane indicates a strong proximal-distal stiffness transition |
| **house sparrow** | **Highest** (23.0 / 29.9) | **Highest** (48.8 / 42.8) | **Lowest** (22.8°) | **Moderate** (1.39; T > L) | **Highest** (4.05×) | Moderate base-to-tip decline in BD (−18% / −21%) and BA (−17% / −19%); barbule density peaks at middle on leading vane; trailing vane decline −12% | Variable: strongly nonlinear for BD, BbD, and barb length asymmetry (edf 5.7-8.3); near-linear for BD asymmetry and BA trailing (edf 1.0); all 11 significant | Reversed barbule asymmetry (0.89; L > T); strongest VW asymmetry (4.05×); elevated BD asymmetry (1.32×) | *Reflects* dense, tightly woven vane structure associated with high-frequency flapping; reversed barbule asymmetry and high VW asymmetry distinctive among species examined |
| **pygmy cormorant** | **Intermediate** (17.8 / 21.5) | **Very low** (22.7 / 24.4) | **Intermediate** (32.4°) | **Low** (1.18; T > L) | **Moderate** (2.46×) | Moderate base-to-tip decrease in BD (leading −27%, trailing −24%) and BA (leading −27%, trailing −22%); barbule density declines distally on both vanes (leading −15%, trailing −22%) | Variable: includes both complex (BbD, BA: edf 5.1-7.5) and near-linear patterns (VW asym: edf 1.0); all 11 significant; lower minimum complexity than WS/CB | Near-symmetric barbule distribution (1.08; T ≈ L); moderate VW asymmetry (2.46×); BD higher on trailing vane (1.23×) | *Reflects* moderately reinforced vane under continuous flapping loads; substantially reduced barbule density (39-53% below other species) may reflect wetttable plumage strategy associated with diving ecology |

### Conclusions

This study provides a detailed multi-scale characterisation of feather vane macrostructure across all remiges in species with contrasting flight styles. Formal comparison of three functional wing regions confirmed that outer primaries possess a distinctive vane macrostructure, with vane width asymmetry two to three times higher than inner primaries or secondaries, reduced leading-vane barb angles, and opposing barb density patterns between vanes. Within-feather longitudinal gradients in barb density and barb angle were conserved across all four species, indicating fundamental structural constraints on force transmission. The interspecific patterns further showed that vane macrostructure might respond to non-flight-related ecological pressures as well: pygmy cormorant’s barbule density was 39–53% lower than in all other species, consistent with its wettable plumage strategy rather than flight demands alone.

Although the small species sample (one per flight style) precludes broad phylogenetic generalisation, the large effect sizes observed for the principal interspecific differences (e.g., 1.3–1.9-fold difference in barb density, 39–53% reduction in barbule density in the pygmy cormorant) indicate these patterns are biologically meaningful and warrant investigation across additional taxa. Future comparative studies combining broad taxonomic coverage with fine-scale spatial resolution and direct mechanical measurements (vane stiffness, air permeability, hydrodynamic resistance) would resolve the relative contributions of phylogeny, body size, flight mechanics, and ecology to vane architecture and test the functional morphology hypotheses proposed here.

The interpretation of feather structural adaptations to different flight styles should be considered as hypothesis based on aerodynamic theory, but not proof for flight-related functional morphology because our four species differ in many biological traits not only in flight style. These interpretations thus should be assessed by future comparative studies with a broader sampling of the avian tree.

## ACKNOWLEDGEMENTS

We thank the members of the Milvus Group Bird and Nature Protection Association and Evolutionary Ecology Group members. Many thanks to Tibor Fuisz, János Déri, Gábor Bakacsi, Andrea Józsa, Ildikó Mező and Márta Osváth-Ferencz for their considerable help in data collection. We are thankful to Mátyás Kassay for the bird and wing drawings.

## DATA AND RESOURCE AVAILABILITY

All data and interactive visualisations are available in the Supplementary Material (Supplementary Material S1: interactive data explorer with raw data visualization, GAM-based estimated marginal means, and feather structure diagrams; Supplementary Material S2: supplementary figures; Supplementary Material S3: supplementary tables).

## LIST OF SYMBOLS AND ABBREVIATIONS

edf: effective degrees of freedom
GAM: generalised additive model
P: primary feather (P1, P2, etc.)
R²adj: adjusted coefficient of determination
S: secondary feather (S1, S2, etc.)
s.e.m.: standard error of the mean

